# The evolutionary cost of homophily: social stratification facilitates stable variant coexistence and increased rates of evolution in host-associated pathogens

**DOI:** 10.1101/2024.07.14.603415

**Authors:** Shuanger Li, Davorka Gulisija, Oana Carja

## Abstract

Coexistence of multiple strains of a pathogen in a host population can present significant challenges to vaccine development or treatment efficacy. Here we discuss a novel mechanism that can increase rates of long-lived strain polymorphism, rooted in the presence of social structure in a host population. We show that social preference of interaction, in conjunction with differences in immunity between host subgroups, can exert varying selection pressure on pathogen strains, creating a balancing mechanism that supports stable viral coexistence, independent of other known mechanisms. We use population genetic models to study rates of pathogen heterozygosity as a function of population size, host population composition, mutant strain fitness differences and host social preferences of interaction. We also show that even small periodic epochs of host population stratification can lead to elevated strain coexistence. These results are robust to varying social preferences of interaction, overall differences in strain fitnesses, and spatial heterogeneity in host population composition. Our results highlight the role of host population social stratification in increasing rates of pathogen strain diversity, with effects that should be considered when designing policies or treatments with a long-term view of curbing pathogen evolution.

## Introduction

From genes to communities, natural systems are characterized by high rates of variant coexistence and understanding the mechanisms that shape and protect the maintenance of diversity in a population is a central problem in biology. Evolutionary mechanisms that are known to promote long-lived, non-neutral strain polymorphism are usually balancing mechanisms, arising from direct negative frequency dependence (Cobey, 2014) or from spatially heterogeneous selection pressure (Lourenço and Recker, 2013). An additional class of mechanisms that have been theoretically shown to promote polymorphism across a wide range of evolutionary scenarios are storage effects, initially recognized in studies of species coexistence (Chesson, 1994, 1997, 2000; Shmida and Ellner, 1984). In systems with spatial, life-stage or genetic background heterogeneity in temporally varying selective pressure, storage effects promote coexistence when a specific life stage a patch of habitat, or genetic background can diminish selection against deleterious variants and store polymorphism in the population until conditions change (Svardal et al., 2011; Svardal and Rueffler, 2015; Gulisija and Kim, 2015; Gulisija et al., 2016). As a result, selection against less fit forms in unfavorable environments is buffered (Chesson, 2000) and intra-species competition is maximized in favorable environments (Ellner and Hairston, 1994; Ellner and Sasaki, 1996; Turelli et al., 2001; Svardal et al., 2011; Svardal and Rueffler, 2015).

Here we ask whether patterns of preferential social interaction between individuals can, similarly, generate enough differential in selection pressure to stably increase rates of variant coexistence in a population. Biased preferences of interaction and their effects on population segregation have been widely documented in the social science literature, starting with the ground-breaking studies of Thomas C. Schelling on how micromotives can shape macro-patterns of population behavior (Schelling, 1978; Centola, 2011; Flatt et al., 2012). Across many dimensions of phenotypes (including physical, cultural, and attitudinal characteristics), humans exhibit high levels of homophily, the tendency to interact with others of similar type, in social tie formation and patterns of social interaction (Sampson et al., 2002; Ioannides and Loury, 2004; McPherson et al., 2001; Kossinets, 2009; Creanza and Feldman, 2014). Moreover, homophily has been shown to significantly shape patterns of pathogen or cultural variant spread. For example, the structure of sexual networks of interaction has been shown to impact the spread of HIV (Janulis et al., 1999) and opinion diffusion through social networks has been shown to influence individuals’ decision for vaccination, thus indirectly also shaping contact-dependent vulnerability to pathogens (Zhou et al., 2015).

In parallel, empirical work has explored how population subgroups of denser interaction also coincide with phenotypic and immunological differences between individuals. Local environmental conditions determined by social interactions have been proposed as key determinants of human cellular immune systems, creating different patterns of interaction between different immunological profiles that could significantly impact pathogen spread and diversity (Leschak and Eisenberger, 2019; Zheng et al., 2020). Vulnerabilities to infections correlate with factors like age, ethnicity, exposure histories, and immune system competence (Zhang et al., 2020b; Bastard et al., 2020; Holmes et al., 2020; Wu and McGoogan, 2020; Fuzele et al., 2020; Verity et al., 2020; Onder et al., 2020; Lau et al., 2020; Davies et al., 2020). Similarly, extensive bacterial strain sharing across human populations, with distinct mother-to-infant, intra-household and intrapopulation transmission patterns, points to the important role of the contact network in shaping microbiome diversity and transmission (Mueller et al., 2015; Song et al., 2013; Brito et al., 2019).

The influence of social patterns of preferential interaction between population groups that are more immunologically and phenotypically similar on the long-term evolutionary dynamics of the pathogen population remains unclear. Here we use a population genetic model to study how social biases of interaction between host population subgroups with varying immune responses or phenotypic characteristics shape rates of pathogen strain polymorphism and long-lived coexistence. We show that, as strains of varying virulence enter the host population, these potential differences in immunity between host subgroups of interaction can promote polymorphism-protecting heterogeneous selective pressure against the pathogen. Therefore, the prevalence of a strain will be increased when circulating within the host group favoring the strain and will be constrained within the social group where the strain is disadvantaged.

This dynamic generates a diversity-promoting mechanism where, unlike typical balancing mechanisms, the diversity is not stored (buffered from selection) in protected life-stages of the pathogen or distinct spatially separated subpopulations, but instead variant diversity is protected by the social interaction structure of the host population. In turn, increased diversity in the pathogen population increases the rate of their evolution, incurring the health cost to the host population, which we refer to as the evolutionary cost of homophily.

We show that increased rates of homophily in the host population promote increased rates of heterozygosity and long-lived multi-strain coexistence and study the role of temporal variance in the host social preference of interaction. We find that periodic amplification of existing homophily in the host population, even when short-term, can significantly increase rates of pathogen diversity and polymorphism.

Our model can also be interpreted through the lens of contagious social behaviors, where two cultural variants are maintained in the population due to the preferential interaction between individuals with similar phenotypic characteristics, such as conformist or anti-conformist behaviors (Muthukrishna et al., 2016; Denton et al., 2020). For both cultural or biological pathogens, understanding the evolutionary mechanisms that contribute to transient or stable variant diversity is essential for designing responses and policies that prevent increases in strain repertoire of pathogenic variants in the population.

### Model description

To examine the effect of social preferences of interaction on rates of host-associated pathogen diversity and polymorphism, we use a Wright-Fisher model to describe changes in strain frequencies within a host population of fixed size *N*. The host population consists of two different types of individuals, with the different host immune-phenotypes denoted by *S* and *A*, interpreted here as individuals with different sensitivity to pathogen virulence, and broadly referred to as symptomatic (*S*) and asymptomatic phenotypes (*A*). These host phenotypic differences correspond to potential differences in immune system composition and effectiveness between people with different genetic backgrounds or in different age groups, which can affect host symptoms and disease progression when infected (Zhang et al., 2020b; Bastard et al., 2020; Holmes et al., 2020; Wu and McGoogan, 2020; Fuzele et al., 2020; Verity et al., 2020; Onder et al., 2020; Lau et al., 2020; Davies et al., 2020). Since the host-population generation times correspond to an equivalent of many microbial generations, we assume that the proportion of *A* to *S* individuals in the host population does not change through the time of pathogen evolutionary dynamics.

Each host individual carries one (and only one) of two different pathogen strains, denoted here by *v/V*. We assume that these two strains differ in their host-pathogen affinity. For example, two different viral strains could differ in their virulence, with the *v* strain having a lower virulence than the *V* strain. Pathogen fitness is determined by the interplay between the virulence of the strain and the immune vulnerability of the host it infects (Fleming-Davies et al., 2018). We model a scenario with trade-off viral fitness distributions on the two host backgrounds *A* and *S*, with optimal virulence smaller on the more sensitive, symptomatic *S* background than on the asymptomatic *A* background (**Figure 1**). It assumes that the benefits of a higher transmission rate can only accrue if the host is healthy enough to interact with other hosts and transmit the strain further. Pathogens with the highest fitness are those with an intermediate level of virulence, striking a balance between within-host production of more transmission forms per unit time and infection strength and length.

**Figure 1:**
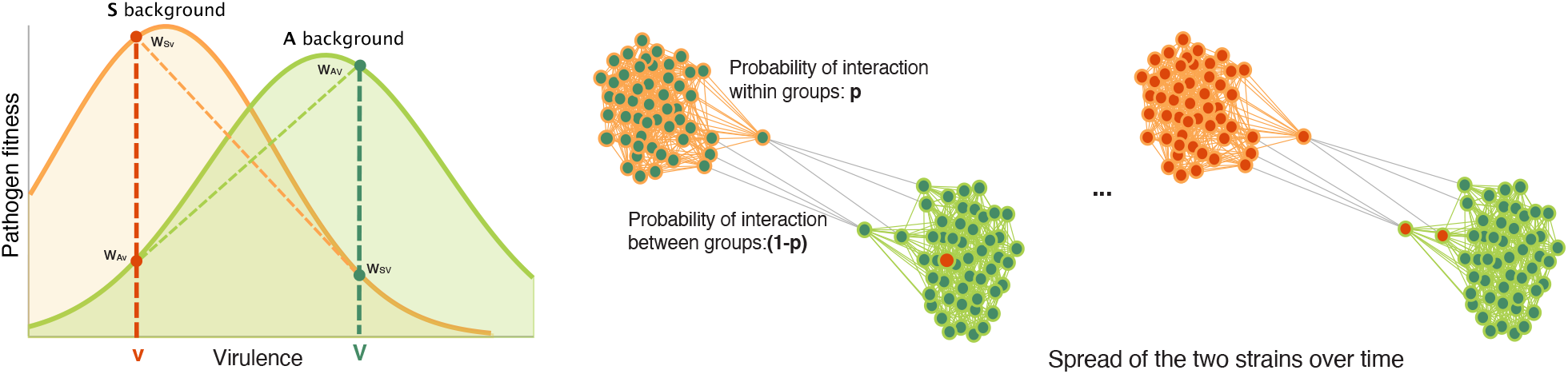
Illustration of the model. Pathogen fitness is determined by the interplay between the virulence of the strain and the immune vulnerability of the host it infects. The more virulent strain *V* is assumed to have higher fitness on the A-type hosts (*w*_*AV*_ *> w*_*Av*_), while strain *v* has higher fitness on the S-type hosts (*w*_*Sv*_ *> w*_*SV*_). The host population is socially structured, with *A* and *S* individuals more likely to interact with other individuals of the same phenotype.

This model can also be applied to understand the evolutionary dynamics of two ‘contagious ‘ cultural traits (for example risk aversion or risk taking behaviors that affect the health of the individual), with different levels of expression on two different individual backgrounds (for example, conformist and anti-conformist hosts or hosts of different status or cultural background (Muthukrishna et al., 2016; Denton et al., 2020)), where trait fitness and transmissibility to other hosts depends on the virulence of the trait, in interplay with its host phenotype and sensitivity to the cultural behavior.

We use the following fitness scheme:

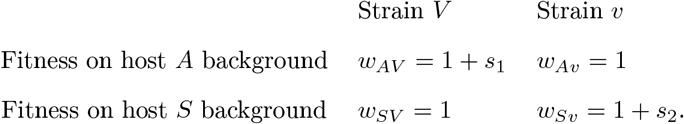

The more virulent strain *V* is assumed to have higher fitness on the *A*-type hosts, while strain *v* has higher fitness on the *S*-type hosts (**Figure 1**). For example, one can interpret the model as symptomatic hosts *S*, infected with a very virulent strain being likely to show symptoms early and reduce social activities, effectively leading to a reduced transmission rate and a selective disadvantage of the *V* strain on this background. On the other hand, immunocompetent hosts *A* show fewer symptoms, even with the *V* strain, which can contribute to higher host transmission rates, thus conferring a selective advantage to the *V* strain on this host background, as has been described in models of viral-host dynamics (McKay et al., 2020). Similarly, for the *v* strain, the lower virulence allows for higher transmission on the symptomatic *S* background. Trade-offs in virulence, transmissibility and pathogen fitness (Anderson and May, 1982; Jensen et al., 2006; Messenger et al., 1999; Mackinnnon et al., 2008; Fraser et al., 2007) are complex and can depend on many parameters. However, the fitness scheme we use is a good baseline model, elegant in its simplicity, to study how the composition and interaction patterns in the host population affect pathogen rates of coexistence.

Since our goal is to study the evolution of the pathogen in the population of infected individuals, we assume that all affected hosts in the population are infected with one of the two strains and, initially, all such hosts are infected with the wild-type strain *v*. The more virulent strain *V* is introduced by a mutation in the host population at the beginning of a simulation run. At each generation, each original host can transmit its pathogen to new hosts. Since a host individual can be infected with a single strain, each subpopulation within a host is considered as an individual unit whose ability to transmit to a next host is determined by strain fitness. Hence, the strain frequencies of the next generation are generated by sampling with replacement proportional to pathogen strain fitnesses and host interaction patterns.

An important aspect of the model is that hosts do not interact randomly, but instead, hosts of the same phenotype are more likely to interact with each other. In other words, there is higher within-group rate of interaction, and hence transmission opportunity, than between-groups (McPherson et al., 2001; Zhang et al., 2020a; Singh and Adhikari, 2020; Centola, 2011; Flatt et al., 2012). We assume a symmetric model where the probability of a host infecting another random host of the same phenotype is given by a fixed preference of interaction, or homophily parameter, *p*, with *p* equal for *A* to *A* and *S* to *S* interactions. Here, *p* represents the probability of infecting a host of the same phenotype. Hosts belonging to different, socially-isolated groups, *A* and *S*, interact with probability (1 − *p*). A population with random interaction groups would have *p* = 0.5.

Strain coexistence and stable polymorphism in the host population can be quantified by the cumulative expected heterozygosity

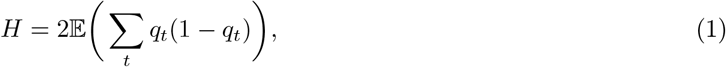

where *q*_*t*_ represents the frequency of strain *V* at time *t* and the expectation is over independent simulations runs. *H* represents the expected sum of heterozygosities over the lifetime of a novel mutant and it can quantify departure from neutrality, independent of population size. When selection is neutral and allele or strain frequencies are only affected by genetic drift, under a randomly mating population, the cumulative expected heterozygosity *H*_*neutral*_ has been shown to be equal to two, regardless of the population size (Kimura, 1969), and thus *H >* 2 describes an elevated level of polymorphism (i.e. coexistence) relative to that under neutrality. It can also be shown that the expected heterozygosity in a haploid population under recurrent mutation *µ* equals *HNµ* (Kimura, 1969).

Social preference of interaction can however generate population subdivision and inflate the cumulative expected heterozygosity *H*. We correct for the effects of population stratification by measuring the subdivided cumulative expected heterozygosity,

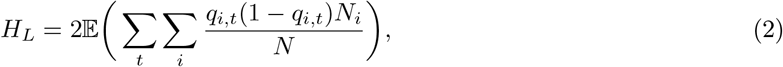

where *q*_*i,t*_ represents the frequency of strain *V* in host social interaction group *i* at time *t*, and *N*_*i*_ represents the size of the host population of immuno-phenotype *i*. The subdivided cumulative expected heterozygosity *H*_*L*_ has also been shown to equal two under neutrality, irrespective of the specifics of population subdivision (Gulisija and Kim, 2015). Therefore, when the levels of polymorphism in the population exceed those under drift, we expect *H*_*L*_ *>* 2 i.e., we observe strain coexistence beyond that expected by chance.

We study the subdivided cumulative expected heterozygosity *H*_*L*_ as a function of the social preference of interaction *p*, the selection coefficients *s*_1_ and *s*_2_, the population size *N*, and the ratio of the two host phenotypes frequencies in the population.

We also extend this baseline model to show its robustness to various relaxations of our assumptions.

#### Temporally varying preference of social interaction

Social preferences of interaction are known to temporally vary (Lofgren et al., 2007). To relax our requirement of constant social interaction preferences, we consider a model in which the hosts change their interaction preferences periodically. We assume hosts interact with preference *p* for *n*_1_ virus generations, and interact without preference for *n*_2_ virus generations. Here, *n*_1_ and *n*_2_ need not be equal.

#### Variance in immune phenotypes within host groups

To reflect the natural demographic variance in host immuno-phenotype and test the model robustness to variance in immune phenotypes within host groups, we develop an extension of the model in which we model host subgroups of fixed population sizes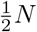, with different proportions of *A* and *S* phenotypes.

#### Changes in host population size and periodic host bottlenecks

Initially, we assume that the infection rates in the host population are such to generate a large number of infected hosts proxied by a constant population size *N*. To accommodate for seasonal changes in infection rates, we also model variable *N*, where the infected population size is initiated at a point in a period of oscillating sizes ranging from 0.05*N* to *N* to 0.05*N*, repeatedly. This generates periods of strong bottlenecks in the affected population.

### Implementation and simulation details

We use Monte Carlo simulations (Gillespie, 1976) to compute the subdivided cumulative expected heterozygosity *H*_*L*_ using an ensemble of at least 5 *×* 10^6^ independent replicate populations of size *N* = 10^5^. In addition to the subdivided cumulative diversity, we record the proportion of simulation runs in which longlived strain polymorphism is maintained in the population at 100N generations. We also record the time of fixation or loss of the two strains, which allows us to quantify the duration of protected polymorphism that perishes. Elevated *H*_*L*_ is compared with values from control simulations either under strain neutrality (all fitness coefficients equal to one) or under a scenario of immune reaction homogeneity across the population.

Simulations are terminated when the new mutant *V* fixes or goes extinct in the population, or, in the case of stable long-lived polymorphism, the two strains coexist over a range of 10^7^ (or 100*N*) generations. We assume a host can only be infected by a pathogen strain obtained from one other host at each generation, there is no co-infection, and the effects of lethality or immunity between generations are negligible.

In the case of temporally varying preference of social interaction, we implement periodic changes in the homophily parameter *p* at deterministic time periods of *n*_1_ and *n*_2_ generations, repeatedly. We implement both symmetric, as well as asymmetric temporal periods, where the duration of the time interval with social preference of interaction can be different from the duration of the time intervals of random interaction and mixing between the hosts.

For the extension of the model to incorporate immune-heterogenous host subgroups, we simulate *R* = 10^7^ independent replicate populations of size *N* = 10^5^ (each host group is assumed to have *N* = 50000). Host phenotype composition are different between the two groups and the probability of *A/A* and *S/S* transmission is now dependent on the *A* and *S* abundances in the two subgroups.

## Results

### Social preferences of interaction promote long-lived pathogen coexistence

Even in a randomly interacting population, the subdivided cumulative expected heterozygosity *H*_*L*_ is a magnitude higher than drift controls, and increases with increasing selective differential *s* (**Figure 2A** and **Supplementary Figure S1**). This is because even though the individuals in the population are randomly interacting, the two host immuno-phenotypes in effect create heterogenous selection, where each host immune-type providing opposite selection pressure on the virus from the other. This protects pathogen diversity in the population for longer than expected under controls.

**Figure 2:**
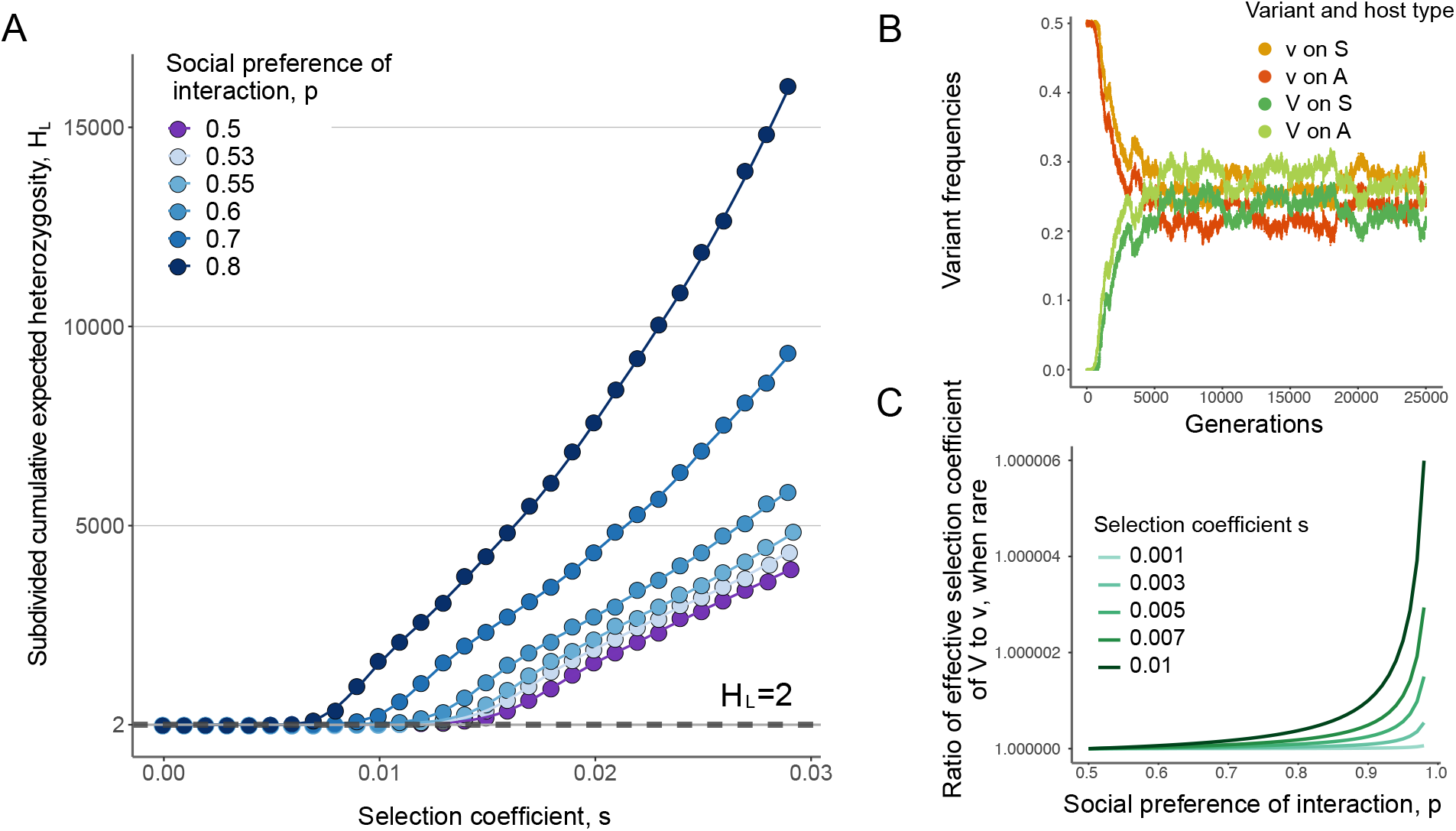
Social preference of interaction promotes increased pathogen strain coexistence. (**A**) The subdivided cumulative expected heterozygosity, *H*_*L*_ in a population of size *N* = 10^5^ as a function of the selection coefficient *s*_1_ = *s*_2_ = *s*. Here, the social preference of interaction *p* is as presented in the legend. The dots represent ensemble averages across 10^7^ replicate Monte Carlo simulations, while the lines represent cubic spline regression. The dotted straight line shows *H*_*neutral*_ = 2. (**B**) Population frequencies of each strain-host combination through time for one run of the simulation, with *N* = 10^5^, *p* = 0.95, and *s*_1_ = *s*_2_ = 0.02. (**C**) Theoretically-derived relative effective selection difference between the two strains when *V* is rare in the population, as a function of the strength of selection and social preference. This effective selection difference increases with both the strength of selection *s* and the social preference of interaction *p*, showcasing the role of negative frequency dependence in shaping patterns of heterozygosity and polymorphism.

As the social stratification of the host population increases (*p >* 0.5), rates of strain heterozygosity raise rapidly, even for small values of social bias of interaction *p* (**Figure 2A**). This rise in levels of cumulative subdivided heterozygosity *H*_*L*_ is more pronounced as the selective differentials between the two fitness strains become stronger. For large enough selection and social preference of interaction, we observe a long-lived polymorphism that stably persists for at least 100*N* generations under a large set of parameter combinations. We sample frequency trajectories to gain insight into the relative evolutionary dynamics between the two strains and find that their frequencies, *f*_*V*_ and *f*_*v*_, oscillate inside a small stable frequency range where *V* preferentially infects *A* hosts and *v* preferentially infects *S* hosts (**Figure 2B**).

Interestingly, even though strains are each preferred on a single host immune type and with strong social stratification (*p* = 0.95), we observe high frequencies of the two strains on both *A* and *S* individuals. In other words, *V* strain proliferation on their preferred *A* hosts, also drive high rates of *V* on *S* hosts in a relatively short period of time, effectively maintaining high frequency of a more virulent strain in the more sensitive host group. Our results are robust to changes in population size *N* and different proportions of *A* to *S* host individuals in the population (**Supplementary Figures S2** and **S3**).

Mechanisms that lead to balancing selection and long-term coexistence can be shown to, directly or indirectly, invoke negative frequency dependence. That is, polymorphism persists if there is a mechanism favoring whichever form is rare. In this model, the increased strain diversity is maintained through indirect negative frequency dependence created by immmune-heterogeneity and social stratification patterns of the host population. Each strain experiences heterogeneous selection through the opposite fitness effects that it experiences on the two different host immune-phenotypes. Here, the opposing selection pressures on the two host immuno-phenotypes generate an association between strain and host that is broken as the pathogen is transmitted to another host. Each strain escapes negative selection in a unfavorable host group through transmission to a favorable host (with probability (1 − *p*)) and increases its numbers by within-group transmission on the preferred host background (with probability *p*) and the combined effect of the two selective forces depends on the frequency of the strain in the population.

To understand the effective negative frequency dependence present in this model, we show that the rare form *V* is indeed favored in a population. Since the two strains have symmetrically opposite fitness effects on two immune phenotypes, the derivation also applies to the *v* strain when rare. Let us assume that the *V* strain appears on its preferred host phenotype *A* and it quickly reaches a type of “interaction-selection balance” as it is transmitted between the two host backgrounds *A* and *S*. At equilibrium, the frequency of *V* on host phenotype *A, x*, is given by

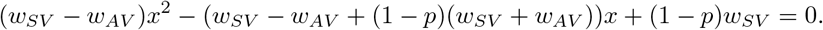

This implies that the equilibrium frequency of phenotype *V* in the *A* host group is given by *f*_*AV*_ :

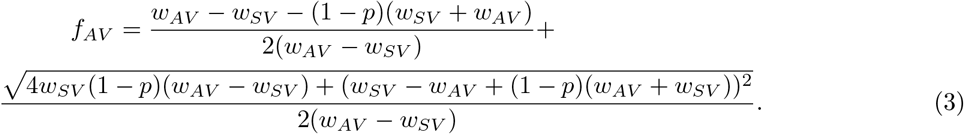

The effective selection coefficient of the *V* strain, when rare, can therefore be written as

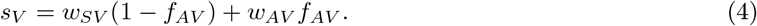

When *s*_*V*_ exceeds the effective selective coefficient of the resident strain *v, s*_*v*_ = (*w*_*Av*_ + *w*_*Sv*_)*/*2 (for equal percentages of *A* and *S* host phenotypes), the *V* mutant strain is preferred, when rare in the population. The effective selection ratio *s*_*effective*_ = *s*_*V,rare*_*/s*_*v,fixed*_ is plotted in **Figure 2C** and shown to increase with both *s* and *p*. Under weak selection, the probability the *V* strain increases past establishment from a single copy in the population can be approximated by twice the effective selection difference between the strains (Haldane, 1927). A similar argument as above reciprocally holds when the strain *v* is rare in a population fixed on the *V* strain.

### The effect of temporally changing social preference of interaction on strain coexistence

Population structure in natural host populations is not constant through time, but can dynamically change, with periods of more population mixing (Lofgren et al., 2007; Zhang et al., 2020a; Singh and Adhikari, 2020). We next ask how elevated rates of polymorphism in the pathogen population change when there are periodic epochs of random and non-random host patterns of interaction. This characteristic also makes our model different from other previous models specifically examining temporally-heterogeneous selection, since changes in social patterns of interaction can act to periodically stratify the population, followed by periods of host random mixing.

We find that social preferences of interaction in the host population need not be constant for elevated pathogen heterozygosity and polymorphism to occur. We consider a periodic temporal regime, with *n*_1_ generations of interaction with fixed preference *p >* 0.5, followed by *n*_2_ generations of random host interaction, *p* = 0.5, which removes population structure. Random host interaction is expected to make intra-strain competition less concentrated in favorable hosts, thereby decreasing heterozygosity. **Figure 3A** shows the values of heterozygosity *H*_*L*_, with *n*_1_ = *n*_2_ = 50 over the whole range of interaction bias. The levels of strain heterozygosity are smaller when host bias changes through time, with inserted epochs of random interaction, but the temporal periods with stratified interactions are sufficient to maintain balanced patterns of pathogen polymorphism (**Supplementary Figure S4**).

**Figure 3:**
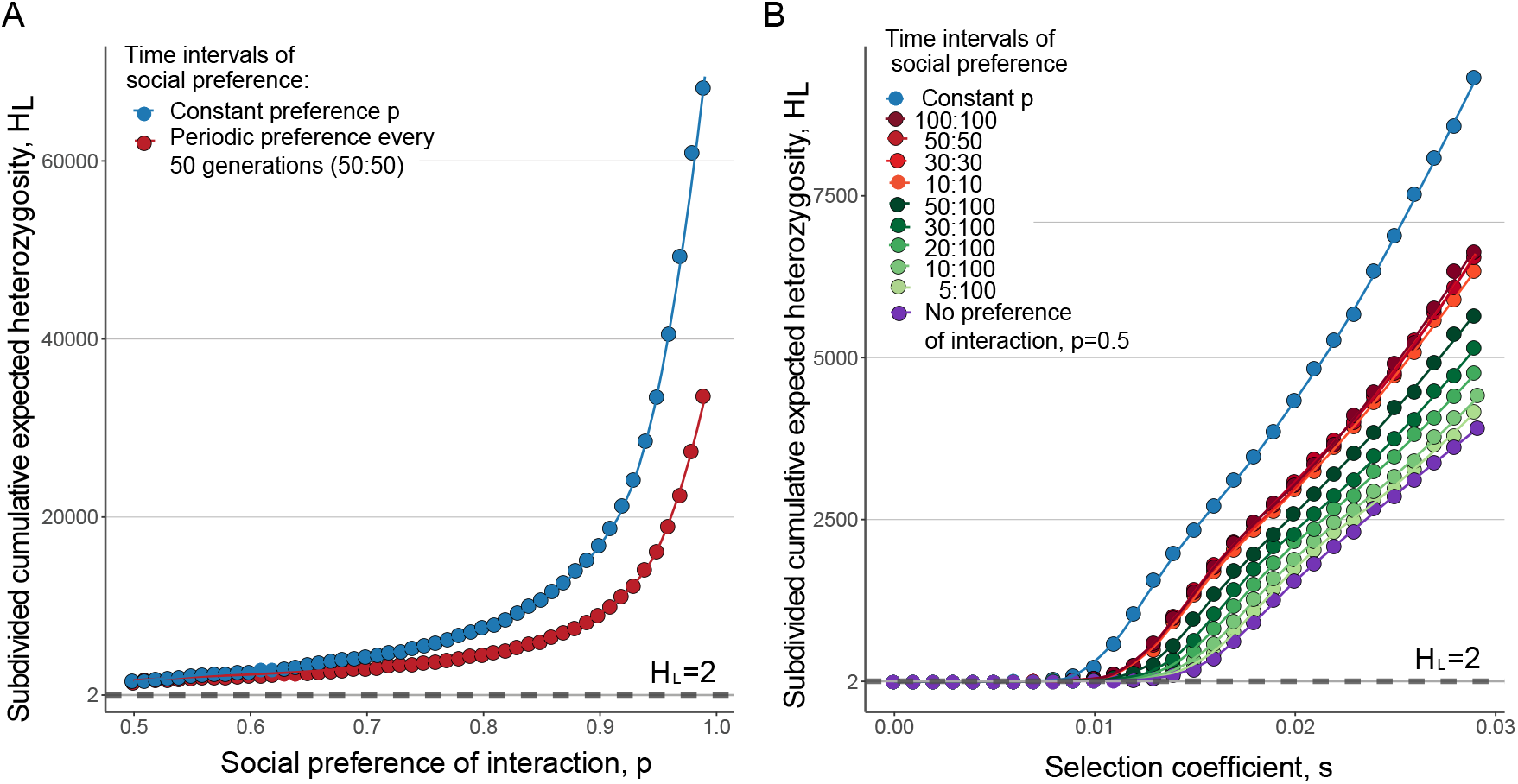
Elevated rates of polymorphism are observed even with periodic interruptions in social patterns of contact. Subdivided cumulative expected heterozygosity, *H*_*L*_ in a population of size *N* = 10^5^ with selection coefficient *s*_1_ = *s*_2_ = *s* = 0.2. The dots represent ensemble averages across 10^7^ replicate Monte Carlo simulations of up to 10^7^ generations, while the lines represent cubic spline regression. (**A**) Comparison of constant preference of interaction *p* with a fluctuating through time regime, where *n*_1_ generations of preference *p >* 0.5 alternate with with *n*_2_ generations with no preference of interaction. Here, *n*_1_ = *n*_2_ = 50. (**B**) Different colors show different *n*_1_, *n*_2_ combinations, with *p* = 0.7.

In fact, even brief periods with non-random host pattern of interaction can promote much higher levels of pathogen coexistence than under random population mixing (**Figure 3B**). We explore different asymmetric combinations of *n*_1_ and *n*_2_ and show, not surprisingly, that smaller 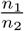. ratios lead to lower levels of balanced polymorphism. Even when bias in interaction is only briefly acting in the population (for example, social interaction preference present a tenth of the time duration), the difference in *H*_*L*_ compared to randomly mixing populations increases with increasing selective differentials between the two mutant strains.

### The effect of strain fitness differences on strain coexistence

Our results are robust across a wide range of parameters and, importantly, they hold in the case of overall fitness differentials between the two strains (*s*_*d*_ = *s*_1_ − *s*_2_ *>* 0). With asymmetric fitness coefficients between the mutants on the two host immuno-phenotypes, the presence of social preference of interaction maintains elevated levels of strain heterozygosity and is expected to be more evident under strong social preference of interaction *p* and small fitness differentials *s*_*d*_. In **Figure 4**, we show a nonlinear trend in the subdivided cumulative expected heterozygosity *H*_*L*_. The strain heterozygosity increases with small differences in the selection coefficients, followed by a decrease as the difference in fitness increases. The larger the selective benefit of the wild-type strain on its preferred host type, the more pronounced this nonlinearity and the increase in population heterozygosity (**Supplementary Figure S5**).

**Figure 4:**
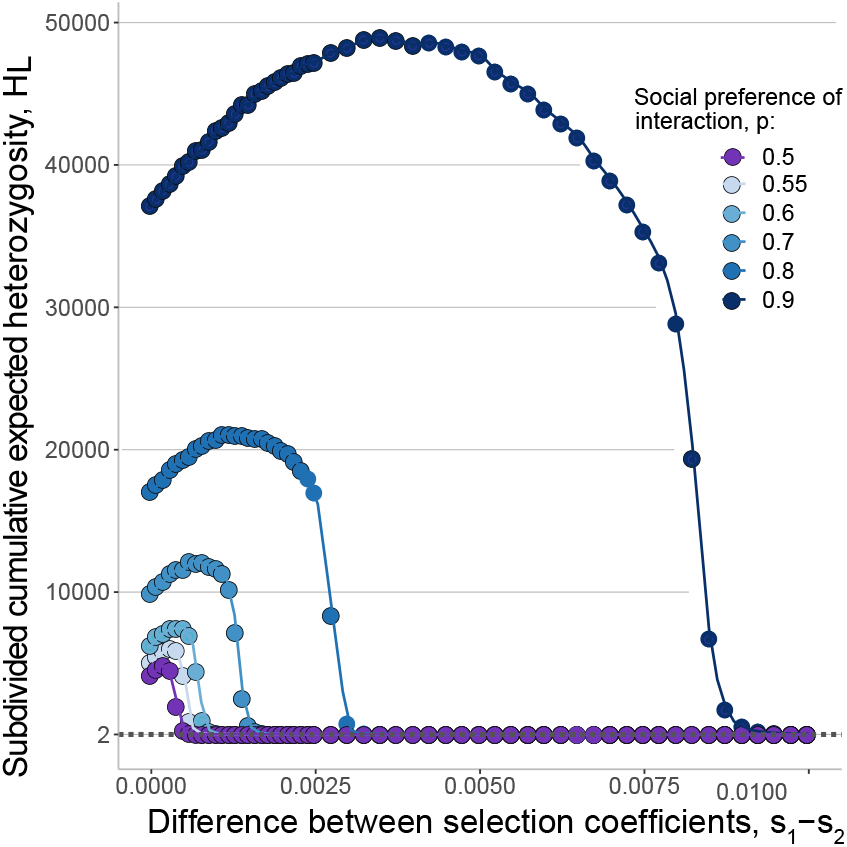
Pathogen strain polymorphism with asymmetric fitness effects between strains. Subdivided cumulative expected heterozygosity, *H*_*L*_ in a population of size *N* = 10^5^ and selection coefficient *s*_2_ = 0.03, 0 ≤ *s*_1_ − *s*_2_ ≤ 0.01. The dots represent ensemble averages across 10^7^ replicate Monte Carlo simulations of up to 10^7^ generations, while the lines represent cubic spline regression.

### The effect of host phenotypic heterogeneity between preferentially interacting individuals

Factors such as exposure histories, age structures, and ethnicities may create immune heterogeneity within socially isolated host groups. We next explore the robustness of the balancing effect arising from our model to immune or phenotypic heterogeneity within host group (*A/S*). We split the population into two host groups and vary the proportion of *A* phenotypes in host subgroup one, *P*_*d*1_(*A*). We set *P*_*d*2_(*A*) = 1 − *P*_*d*1_(*A*), with individuals interacting randomly within the two groups. The strain balancing effect of heterogenous selection and social distancing is robust to within-group immune diversity and is greater with greater differences in the proportion of *A/S* within groups (**Figure 5**). However, coexistence occurs even when the two groups have the same proportions of immuno-phenotypes and are not socially distanced. As previously observed, the effect increases with *p*.

**Figure 5:**
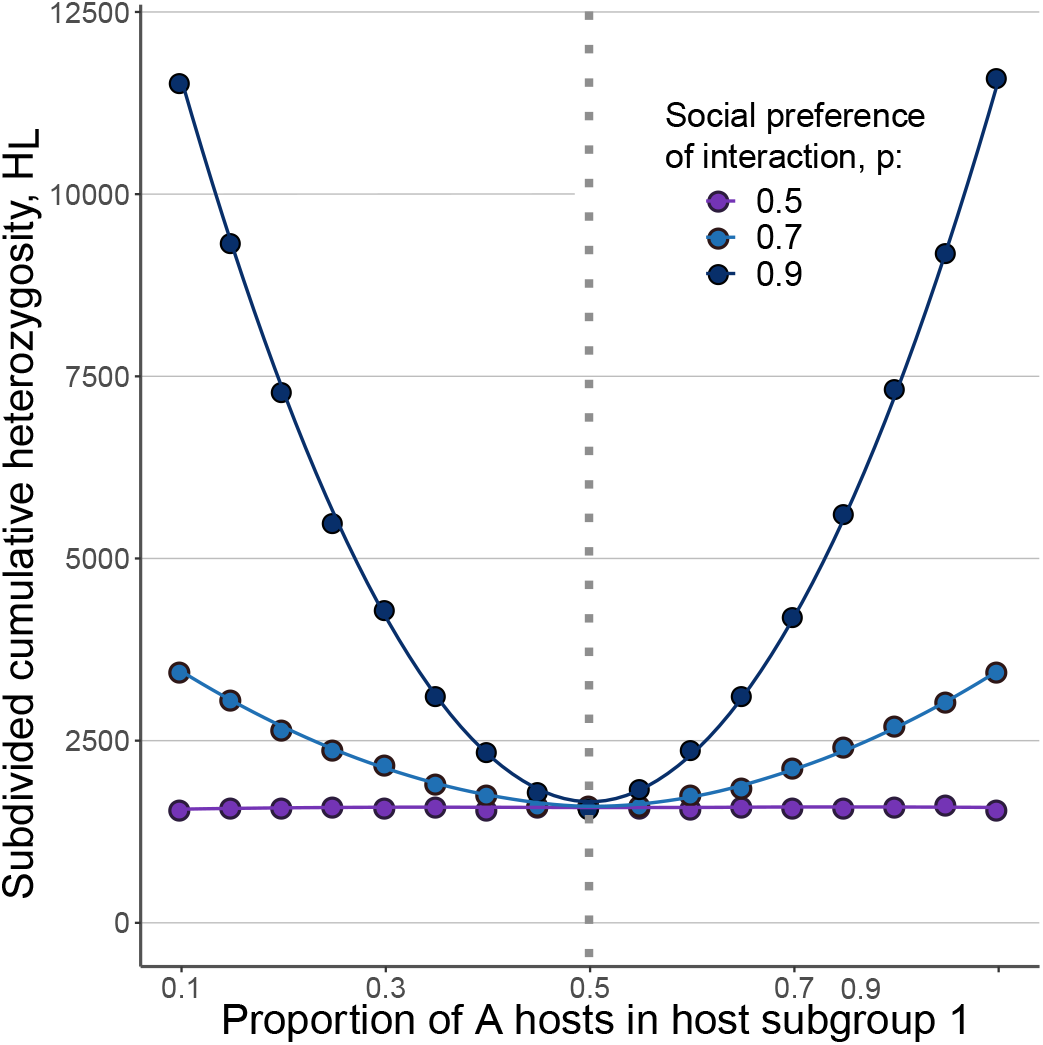
Strain coexistence with heterogeneous host population subgroups. The dots represent ensemble averages across 10^7^ replicate Monte Carlo simulations of up to 10^7^ generations, while the lines represent cubic spline regression. Here, *N*_1_ = *N*_2_ = 5 *×* 10^4^, *s*_1_ = *s*_2_ = 0.02 and random interaction within the subgroups but biased interaction between subgroups, as in legend. The proportion of *A* immune-phenotypes in group one is given on the *x* axis.

## Discussion

As populations continuously adapt from one environment to the other, genetic polymorphism provides a readily available reservoir of adaptive alleles that selection can act upon and thus promotes population persistence (Lande and Shannon, 1996; Barrett and Schluter, 2008; Pennings, 2012; Furuse and Oshitani, 2016). Therefore coexistence of multiple pathogen strains in a host population poses significant challenges for vaccine development or treatment outcome. For example, if a vaccine only confers immunity to a subset of viruses circulating in the population, other variants can persist and become reservoirs for viral evolution and further vaccine evasion (Yen and Webster, 2009; Carrat and Flahault, 2007; Furuse and Oshitani, 2016). A central question thus becomes: which mechanisms enable multiple strain coexistence? Here we show how social stratification, combined with the existence of distinct host immuno-phenotypes, can lead to increased levels of pathogen strain or contagious cultural behavior coexistence and highlight the importance of social preference of interaction in promoting long-lived pathogen polymorphism and rates of evolution.

Mechanisms that promote long-term coexistence and maintain balanced polymorphism in a population, can be shown to, directly or indirectly, invoke some kind of negative frequency dependence. Direct negative frequency dependence acts through the interaction of host immunity and virus immunophenotypes, whereby common strains experience more intra-host competition due to prior host exposure to similar strains (Cobey, 2014). This favors the evolution of antigenically diverse strains, as seen in influenza, rotavius, and HPV (Zinder et al., 2017a,b; Ranjeva et al., 2017).

Here, we discuss a mechanism that promotes multi-strain pathogen coexistence through indirect negative frequency dependence driven by elevated biases of interaction between individuals in distinct immunophenotype groups in the host population. The complex interactions of human society result in heterogenous contact networks where interactions are common between some individuals in a population and are entirely absent between others (Christakis and Fowler, 2008, 2013; Perc et al., 2013). This stratification and sparsity of connections in human populations has previously been shown to affect the culture-wide adoptions of behaviors that can have different effects on fitness in different environments or individual phenotypic backgrounds (Carja and Creanza, 2019). Similarly, geographic clustering of a host population, which promotes spatio-temporal structures with varying immune selection, has been shown to affect viral diversity and evolution (Holmes, 2004; Lourenço and Recker, 2013).

The mechanism of coexistence in our model is rooted in the fact that different strains of the pathogen experience different selection pressures when infecting hosts of different immune vulnerability, i.e. immunophenotypes. Since host interaction patterns can also coincide with host vulnerability, prevalence of a strain is expected to be promoted within the favorable interaction group. Because host populations are immune-heterogeneous and hosts from different groups also interact, strain prevalence is also influenced by inter-strain competitions.

We show that long-lived strain coexistence is promoted by the immune heterogeneity between hosts and the magnitude of host population stratification. Our results are robust to overall fitness differences between the two mutant strains, changes in population size *N* and different proportions of *A* to *S* host individuals in the population, as well as temporal fluctuations in interaction patterns and within-group heterogeneity. We show that temporal fluctuations in the strength of the social preference of host interaction reduce rates of polymorphism, but stratification nonetheless is the dominant effect, maintaining elevated levels of heterozygosity in the pathogen population.

It is important to note that our mechanism alone is fast enough to create multi-strain coexistence in short time scales (**Supplementary Figures S6**). We show that strain coexistence is achieved in a much shorter period of time in a stratified host population compared to a randomly interacting one, and it is maintained for long time scales. These results imply high evolvability in new pathogens with high transmission rates that are constrained between social groups, but allowed to spread within groups.

As a proof-of-concept, our model assumes constant proportions of *A* and *S* hosts in a population. While our robustness analysis shows that the effect we report holds across various proportions of *A* and *S* within preferentially interacting groups, we do not test temporal variation in their ratio that might arise due to immigration of hosts to the population, for example. Likewise, temporal variations in interaction preferences should consider non-deterministic patterns and, to better capture multi-strain coexistence, this model could be extended to multiple strains and multiple host groups. Further extensions of the model could also take into account other types of host-pathogen interactions. For example, different strains of a viral population can infect the same host, resulting in competition or facilitation (Rousseau et al., 2001; Mavilia and Wu, 2018). Development of acquired immunity should also be considered, especially vulnerability of re-infection (Gousseff et al., 2020; Parry, 2020). Additionally, as multi-strain coexistence is promoted by multiple mechanisms (Cobey, 2014; Zinder et al., 2017a,b; Ranjeva et al., 2017; Lourenço and Recker, 2013; Holmes, 2004), it is important to consider the interplay of these mechanisms when making evolutionary predictions.

Lastly, while we find that the reported effect is robust to changes in the infection rates in the host population, by effectively modeling severe oscillating bottlenecks in the number of affected individuals (**Supplementary Figure S7**), we assume that the resident strain is already circulating in the population at the time an invader strain is introduced. While experiencing small infection rate (bottleneck) could serve as a proxy for a new outbreak, we did not model the outbreak itself and further models could accommodate for a wider set of epidemiological scenarios.

## Acknowledgments

This research was done using resources provided by the Open Science Grid, which is supported by the National Science Foundation award 1148698, and the U.S. Department of Energy’s Office of Science.

## Funding

We gratefully acknowledge support from the NIH National Institute of General Medical Sciences (award no. R35GM147445) and the United States-Israel Binational Science Foundation (award no. 2019266).

## Competing interests

There are no competing interests.

## Supplementary Figures

**Figure S1:**
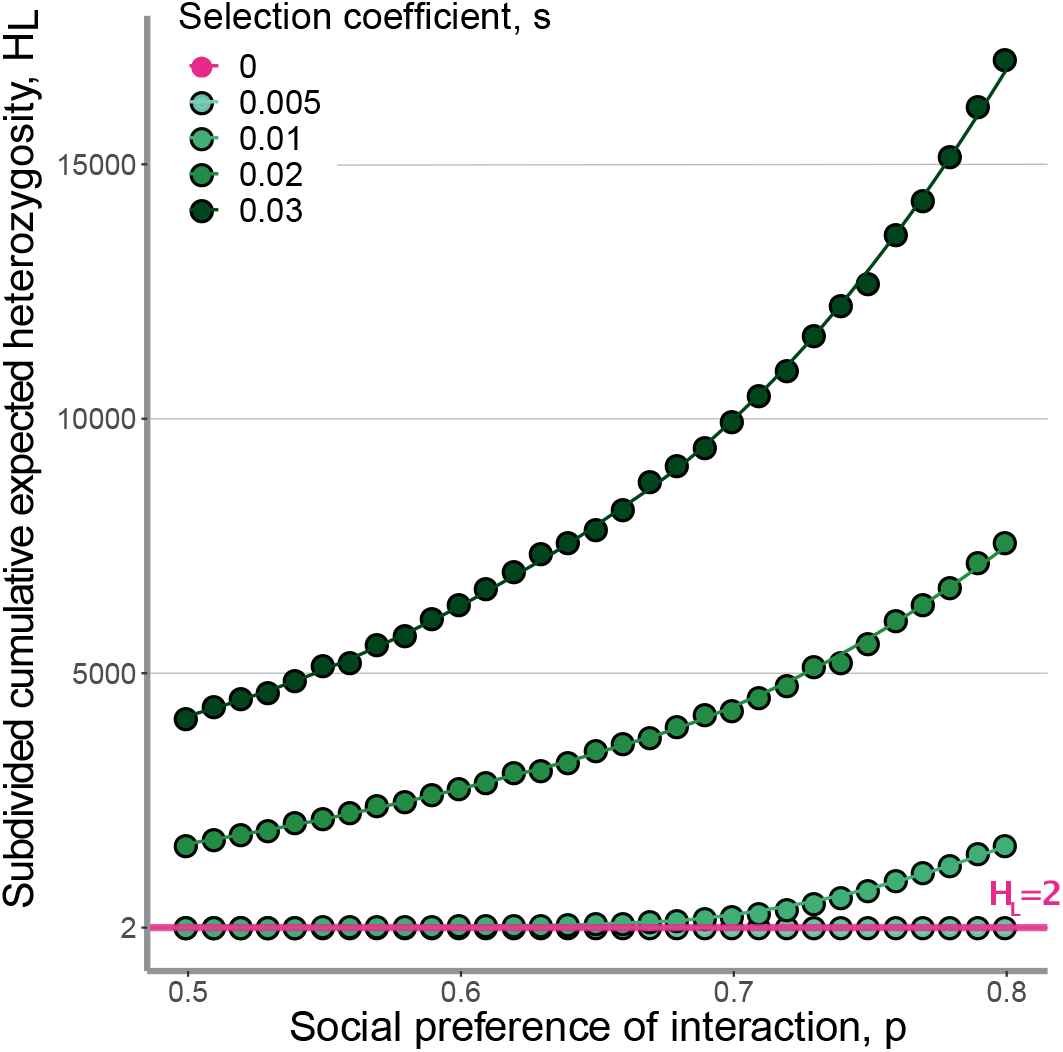
Social preference of interaction promotes increased rates of strain heterozygosity and coexistence. The role of symmetric selection coefficient *s*. The dots represent ensemble averages across 10^7^ replicate Monte Carlo simulations of up to 10^7^ generations, while the lines represent cubic spline regression. Subdivided cumulative expected heterozygosity, *H*_*L*_ in a population of size *N* = 10^5^ with social preference of interaction *p* as on the *x* axis. The colors represent different values of selection coefficient, *s*_1_ = *s*_2_ = (0, 0.005, 0.01, 0.02, 0.03), as presented in the legend. We show that *H*_*L*_ increases as *p* and *s*_1_ = *s*_2_ = *s* increase.

**Figure S2:**
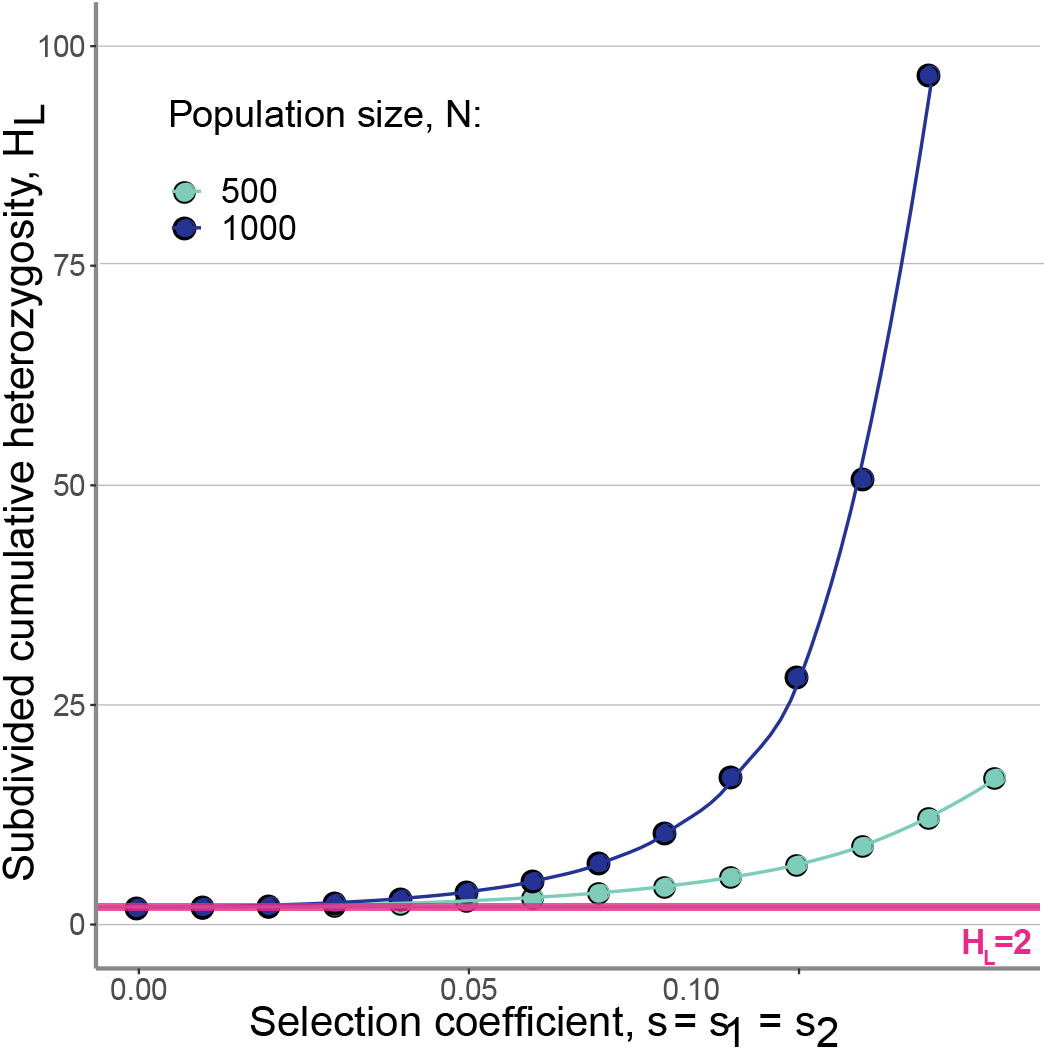
Social preference of interaction promotes elevated polymorphism even when population size is small. The dots represent ensemble averages across 10^7^ replicate Monte Carlo simulations of up to 10^7^ generations, while the lines represent cubic spline regression. Subdivided cumulative expected heterozygosity, *H*_*L*_, in a relatively small population, with social preference of interaction *p* = 0.6. The colors represent different population sizes *N* = (500, 1000).

**Figure S3:**
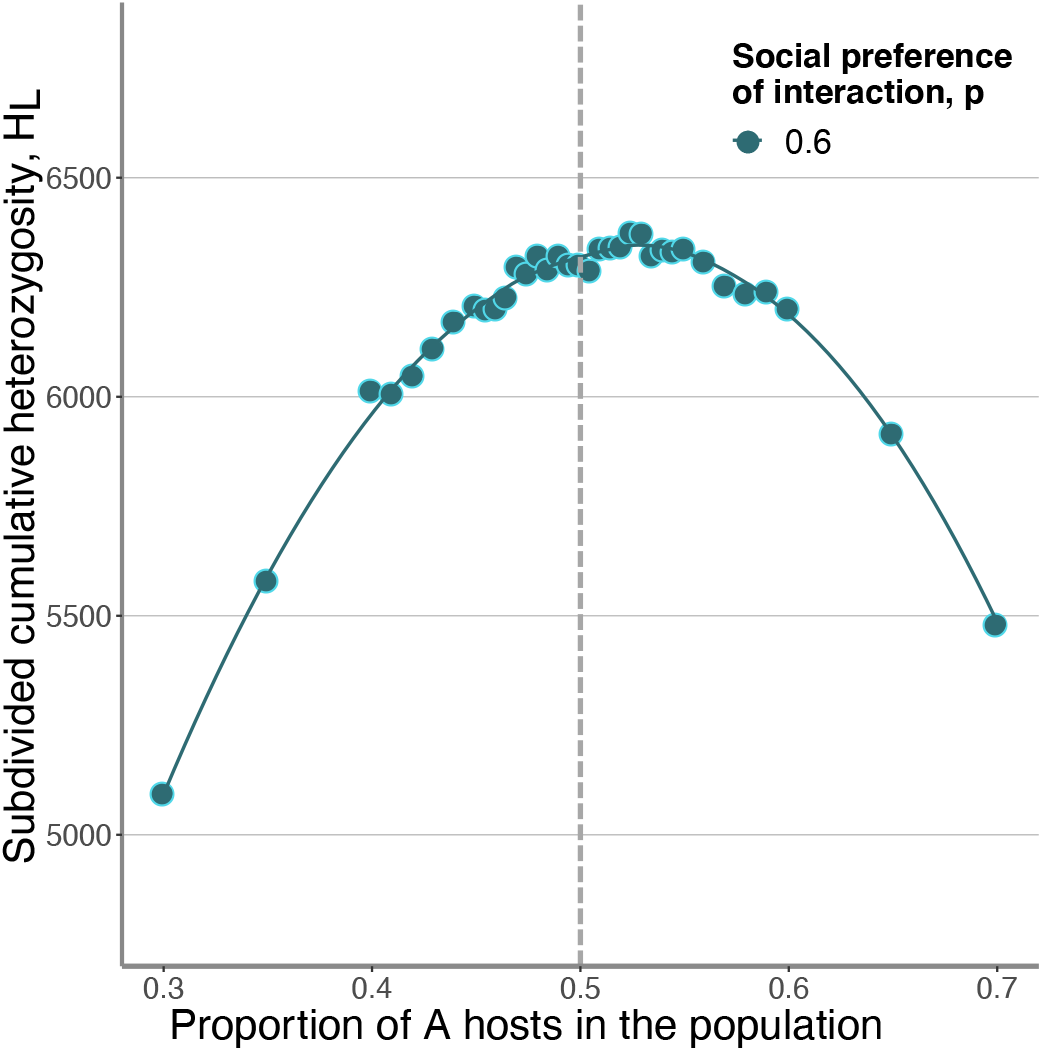
Levels of heterozygosity as a function of *A* and *S* proportions in the host population. The dots represent ensemble averages across 10^7^ replicate Monte Carlo simulations of up to 10^7^ generations, while the lines represent cubic spline regression. Subdivided cumulative expected heterozygosity, *H*_*L*_ for various frequencies of *A* phenotype in the host population as presented on the *x* axis, with *N* = 10^5^, selection coefficients *s*_1_ = *s*_2_ = 0.03, and social preference of interaction *p* = 0.6.

**Figure S4:**
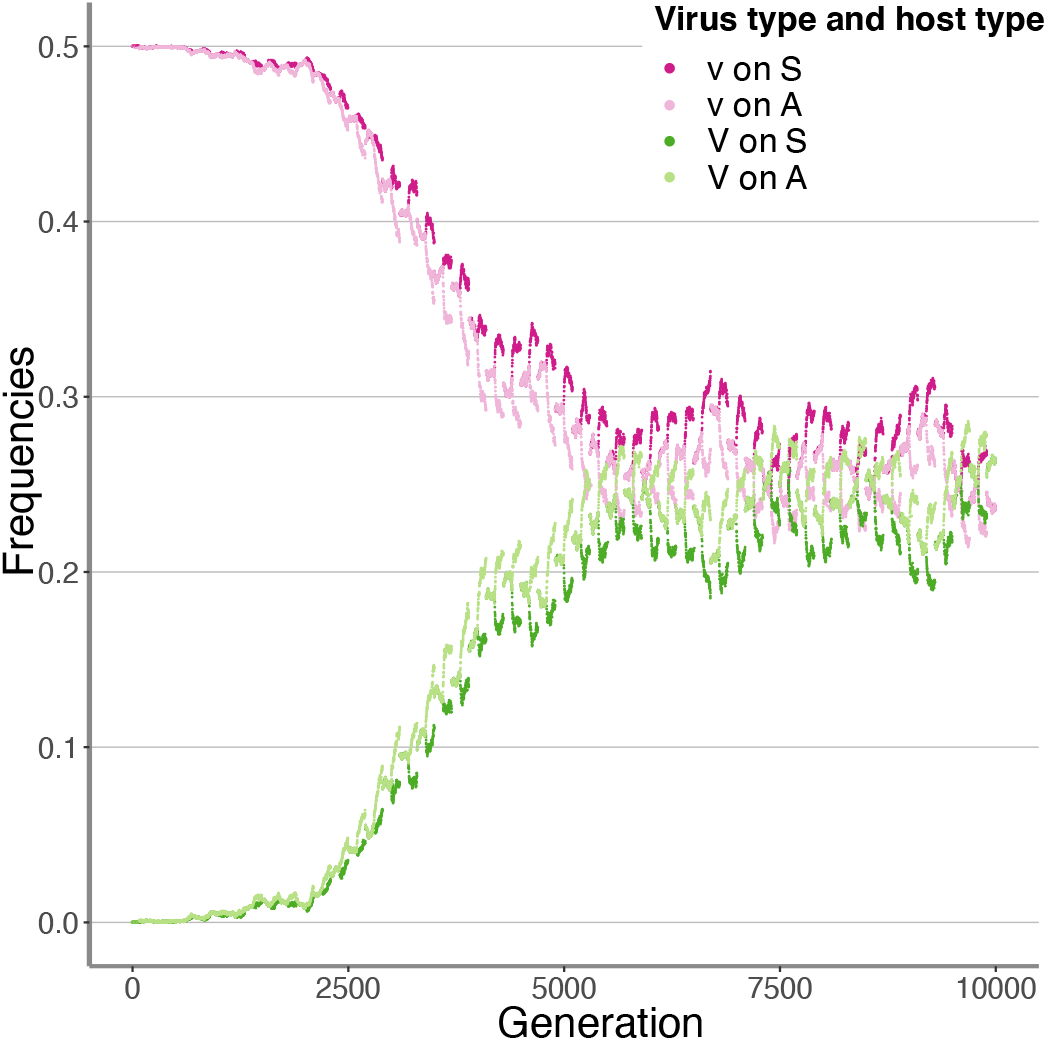
Periodic presence of preference promotes pathogen strain coexistence. Simulated population frequencies of each strain-host combination through time for one simulation run. Here, *N* = 10^5^, *s*1 = *s*2 = 0.02. Periods of preference (*n*_1_) switch with periods of no preference (*n*_2_), with *n*_1_ = *n*_2_ = 100.

**Figure S5:**
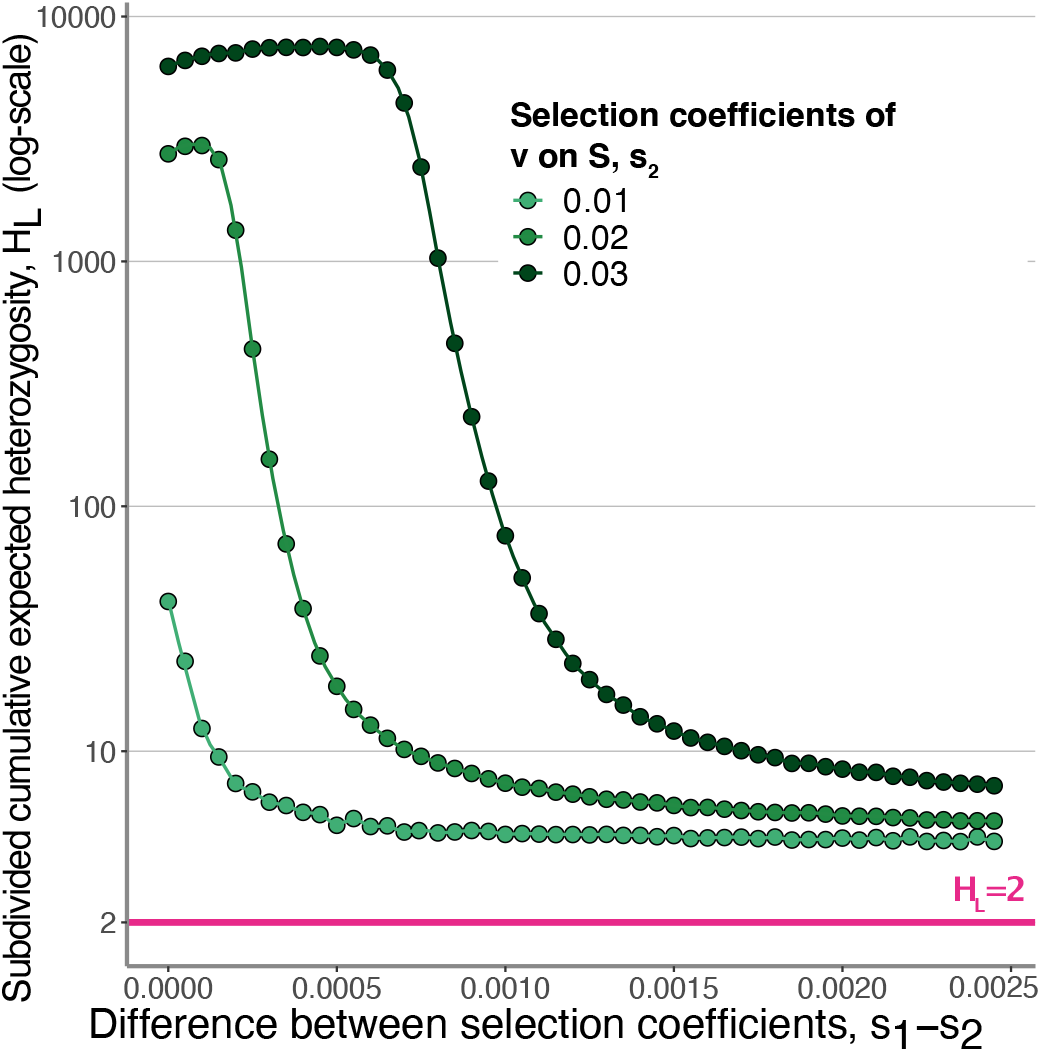
Effect of selection coefficient on strain polymorphism when the two mutant strains have overall difference in fitness. The dots represent ensemble averages across 10^7^ replicate Monte Carlo simulations of up to 10^7^ generations, while the lines represent cubic spline regression. Subdivided cumulative expected heterozygosity, *H*_*L*_ in a population of size *N* = 10^5^, 0 ≤ *s*_1_ − *s*_2_ ≤ 0.0025, and social preference of interaction *p* = 0.6. The different colors represent *s*_2_ = (0.01, 0.02, 0.03).

**Figure S6:**
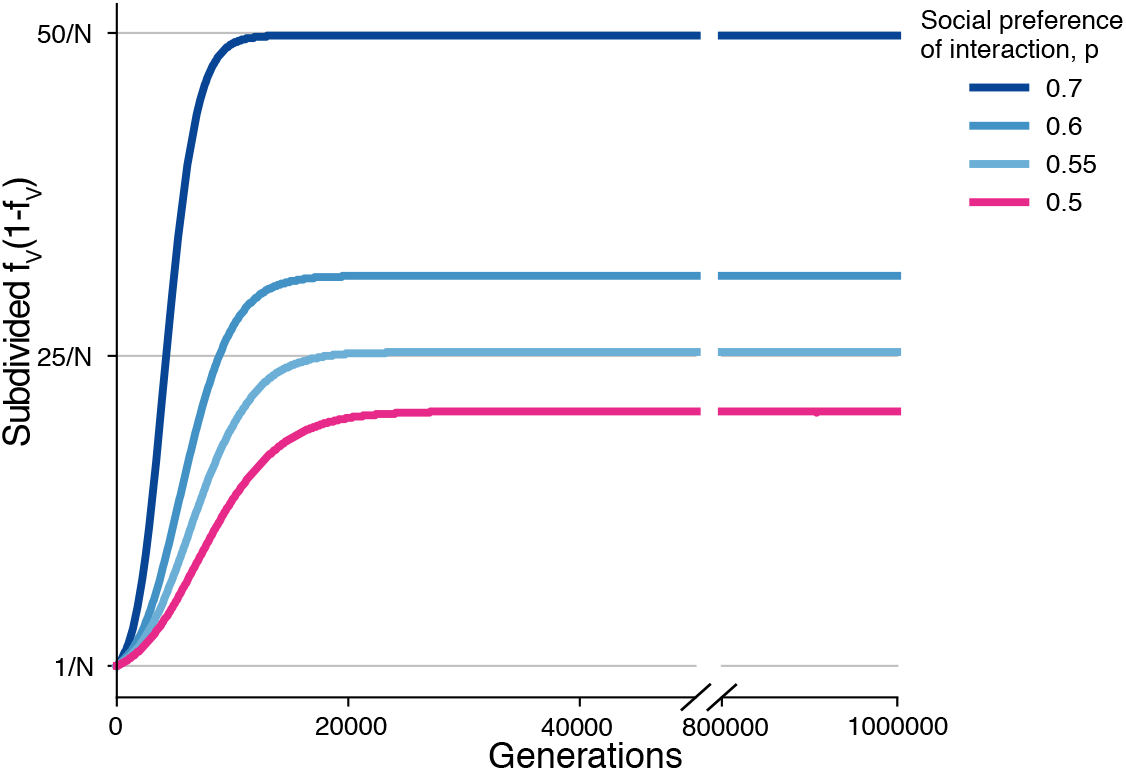
Fast multi-strain coexistence in stratified host populations. Averaged simulation results through time. Here, selection coefficient *s*_1_ = *s*_2_ = 0.03 in a population of *N* = 10^5^, with 10^5^ repetitions. *f*_*V*_ represents frequency of *V* on each of the host backgrounds, *A* and *S*. Subdivided *f*_*V*_ *×* (1 − *f*_*V*_) measures the sum of *f*_*V*_ *×* (1 − *f*_*V*_) weighted by the host population sizes: 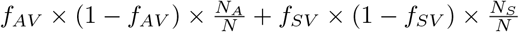.

**Figure S7:**
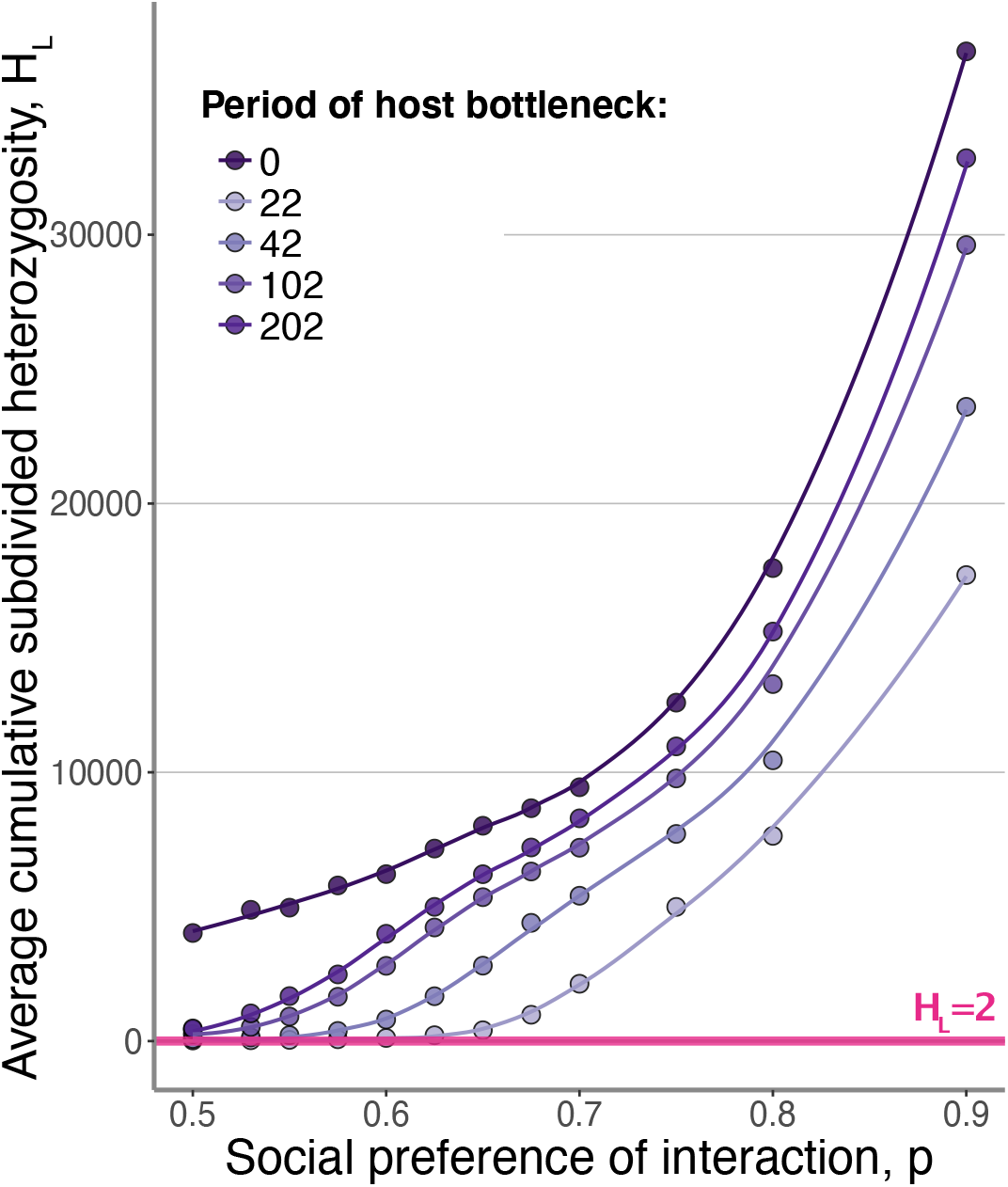
Robustness of the polymorphism promoting effect by homophily to periodic changes in affected population size, including population bottlenecks.. Here, we performed 100*N* replicate simulation runs assuming an oscillating population size ranging from 0.05*N* to *N* repeatedly, starting at a random point in this cycle. The ratio *S* to *A* is 0, *s*_1_ = *s*_2_ = 0.03 and *N* = 100000.

## Notes

### Competing Interest Statement

The authors have declared no competing interest.

## References

Roy M. Anderson and Robert M. May. Coevolution of hosts and parasites. Parasitology, 84:411–426, 1982.

Rowan D. H. Barrett and D. Schluter. Adaptation from standing genetic variation. Trends in Ecology & Evolution, 23(1):38–44, 2008.

Paul Bastard, Lindsey B. Rosen, Qian Zhang, et al. Autoantibodies against type I IFNs in patients with life-threatening COVID-19. Science, 370(6515), 2020.

Ilana L Brito, Thomas Gurry, Shijie Zhao, Katherine Huang, Sarah K Young, Terrence P Shea, Waisea Naisilisili, Aaron P Jenkins, Stacy D Jupiter, Dirk Gevers, et al. Transmission of human-associated microbiota along family and social networks. Nature Microbiology, 4(6):964–971, 2019.

Oana Carja and Nicole Creanza. The evolutionary advantage of cultural memory on heterogeneous contact networks. Theor Popul Biol, 129:118–125, 2019.

Fabrice Carrat and Antoine Flahault. Influenza vaccine: The challenge of antigenic drift. Vaccine, 25(39): 6852–6862, 2007.

Damon Centola. An experimental study of homophily in the adoption of health behavior. Science, 334 (6060):1269–1272, 2011.

Peter Chesson. Multispecies competition in variable environments. Theoretical Population Biology, 45:227–276, 1994.

Peter Chesson. Diversity maintenance by integration of mechanisms over various scales. Proc. 8th International Coral Reef Symposium, 1:405–410, 1997.

Peter Chesson. Mechanisms of maintenance of species diversity. Annual Review of Ecology and Systematics, 31(1):343–366, 2000.

NA Christakis and JH Fowler. The collective dynamics of smoking in a large social network. N Engl J Med, 358(21):2249–58, 2008.

NA Christakis and JH Fowler. Social contagion theory: examining dynamic social networks and human behavior. Stat Med, 32:556–577, 2013.

Sarah Cobey. Pathogen evolution and the immunological niche. Annals of the New York Academy of Sciences, 1320(1):1–15, 2014.

Nicole Creanza and Marcus W. Feldman. Complexity in models of cultural niche construction with selection and homophily. Proceedings of the National Academy of Sciences, 111(Supplement 3):10830–10837, 2014.

Nicholas G. Davies, Petra Klepac, Yang Liu, Kiesha Prem, and Mark Jit. Age-dependent effects in the transmission and control of COVID-19 epidemics. medRxiv, 2020.

Kaleda Krebs Denton, Yoav Ram, Uri Liberman, and Marcus W. Feldman. Cultural evolution of conformity and anticonformity. Proceedings of the National Academy of Sciences, 117(24):13603–13614, 2020.

Stephan Ellner and Nelson G. Hairston. Role of overlapping generations in maintaining genetic variation in a fluctuating environment. The American Naturalist, 143(3):403–417, 1994.

Stephan Ellner and Akira Sasaki. Patterns of genetic polymorphism maintained by fluctuating selection with overlapping generations. Theoretical Population Biology, 50(1):31–65, 1996.

Jason D. Flatt, Yll Agimi, and Steve M. Albert. Homophily and health behavior in social networks of older adults. Familiy and Community Health, 35(4):312–321, 2012.

Arietta E. Fleming-Davies, Paul D. Williams André, A. Dhondt, et al. Incomplete host immunity favors the evolution of virulence in an emergent pathogen. Science, 359(6379):1030–1033, 2018.

Christophe Fraser, Déirdre Hollingsworth, Ruth Chapman, Frank de Wolf, and William P. Hanage. Variation in HIV-1 set-point viral load: epidemiological analysis and an evolutionary hypothesis. PNAS, 104(44):17441–17446, 2007.

Yuki Furuse and Hitoshi Oshitani. Mechanisms of replacement of circulating viruses by seasonal and pandemic influenza A viruses. International Journal of Infectious Diseases, 51:6–14, 2016.

Sadanand Fuzele, Bikash Sahay, Ibrahim Yusufu, Tae Jin Lee, Ashok Sharma, Ravindra Kolhe, and Isalesm Carlos M. COVID-19 virulence in aged patients might be impacted by the host cellular MicroRNAs abundance/profile. Aging and Disease, 11(3):509–522, 2020.

Daniel T. Gillespie. A general method for numerically simulating the stochastic time evolution of coupled chemical reactions. Trends in Ecology & Evolution, 22:403–434, 1976.

Marie Gousseff, Pauline Penot, Laure Gallay, et al. Clinical recurrences of COVID-19 symptoms after recovery: Viral relapse, reinfection or inflammatory rebound? Journal of Infection, 81:816–846, 2020.

Davorka Gulisija and Yuseob Kim. Emergence of long-term balanced polymorphism under cyclic selection of spatially variable magnitude. Evolution, 69(4):979–992, 2015.

Davorka Gulisija, Yuseob Kim, and Joshua B. Plotkin. Phenotypic plasticity promotes balanced polymorphism in periodic environments by a genomic storage effect. Genetics, 202(4):1437–1448, 2016.

J. B. S. Haldane. A mathematical theory of natural and artificial selection, part V: Selection and mutation. Mathematical Proceedings of the Cambridge Philosophical Society, 23(7):838–844, 1927.

Edward C. Holmes. The phylogeography of human viruses. Molecular Ecology, 13(4):745–756, 2004.

Laurens Holmes, Michael Enwere, Janille Williams, et al. Black-White risk differentials in COVID-19 (SARS-COV2) transmission, mortality and case fatality in the United States: Translational epidemiologic perspective and challenges. International Journal of Environmental Research and Public Health, 17(12):4322, 2020.

Yannis M. Ioannides and Linda D. Loury. Job information networks, neighborhood effects, and inequality. Journal of Economic Literature, 42(4):105–1093, 2004.

Patrick Janulis, Gregory Phillips, Michelle Birkett, and Brian Mustanski. Sexual networks of racially diverse young MSM differ in racial homophily but not concurrency. Journal of acquired immune deficiency syndromes, 77(5):459–466, 1999.

Knut H. Jensen, Tom Little, Arne Skorping, and Dieter Ebert. Empirical support for optimal virulence in a castrating parasite. PLOS Biology, 4(7):e197, 2006.

Motoo Kimura. The number of heterozygous nucleotide sites maintained in a finite population due to steady flux of mutations. Genetics, 61(4):893–903, 1969.

Gueorgi Kossinets. Origins of homophily in an evolving social network. American Journal of Sociology, 115 (2):459–466, 2009.

Russell Lande and Susan Shannon. The role of genetic variation in adaptation and population persistence in a changing environment. Evolution, 50(1):434–437, 1996.

Max S.Y. Lau, Bryan Grenfell, Kristin Nelson, and Ben Lopman. Characterizing super-spreading events and age-specific infectiousness of SARS-CoV-2 transmission in Georgia, USA. PNAS, 36(117):22430–22435, 2020.

Carrianne J Leschak and Naomi I Eisenberger. Two distinct immune pathways linking social relationships with health: inflammatory and antiviral processes. Psychosomatic Medicine, 81(8):711, 2019.

Eric Lofgren, Nina H. Fefferman, Yuri N. Naumov, Jack Gorski, and Elena N. Naumova. Influenza seasonality: Underlying causes and modeling theories. Journal of Virology, 81(11):5429–5436, 2007.

José Lourenço and Mario Recker. Natural, persistent oscillations in a spatial multi-strain disease system with application to dengue. PLOS Computational Biology, 9(10):e1003308, 2013.

Margaret J. Mackinnnon, Sylvain Gandon, and Andrew Read. Virulence evolution in response to vaccination: The case of malaria. Vaccine, 26:C42–C52, 2008.

Marianna G. Mavilia and George Y. Wu. HBV-HCV coinfection: Viral interactions, management, and viral reactivation. Journal of Clinical and Translational Hepatology, 6(3):296–305, 2018.

Brian McKay, Mark Ebell, Ariella Perry Dale, Ye Shen, and Andreas Handel. Virulence-mediated infectiousness and activity trade-offs and their impact on transmission potential of influenza patients. PNAS, 287 (1927):2020049, 2020.

Miller McPherson, Lynn Smith-Lovin, and James M. Cook. Birds of a feather: Homophily in social networks. Annual Review of Sociology, 27(1):415–444, 2001.

Sharon L. Messenger, Ian J. Molineux, and Bull J. J. Virulence evolution in a virus obeys a trade-off. Proceedings of the Royal Society of London. Series B: Biological Sciences, 266(1417):397–404, 1999.

Noel T Mueller, Elizabeth Bakacs, Joan Combellick, Zoya Grigoryan, and Maria G Dominguez-Bello. The infant microbiome development: mom matters. Trends in Molecular Medicine, 21(2):109–117, 2015.

M. Muthukrishna, T.J. Morgan, and J. Henrich. The when and who of social learning and conformist transmission. Evol. Hum. Behav., pages 10–20, 2016.

Graziano Onder, Giovanni Rezza, and Silvio Brusaferro. Case-fatality rate and characteristics of patients dying in relation to COVID-19 in Italy. JAMA, 2020.

Jane Parry. Covid-19: Hong Kong scientists report first confirmed case of reinfection. BMJ, 370, 2020.

Pleuni Simone Pennings. Standing genetic variation and the evolution of drug resistance in HIV. PLOS Computational Biology, 8(6):e1002527, 2012.

M Perc, J Gomez-Gardenes, A Szolnoki, LM Floria, and Y Moreno. Evolutionary dynamics of group interactions on structured populations: a review. J R Soc Interface, 10: 20120997, 2013.

Sylvia L. Ranjeva, Edward B. Baskerville, Vanja Dukic, Luisa L. Villa, Eduardo Lazcano-Ponce, Ann R. Giuliano, Greg Dwyer, and Sarah Cobey. Recurring infection with ecologically distinct HPV types can explain high prevalence and diversity. PNAS, 114(51):13573–13578, 2017.

Marie-Claude Rousseau, Joao S. Pereira, et al. Cervical coinfection with human papillomavirus (HPV) types as a predictor of acquisition and persistence of HPV infection. The Journal of Infectious Diseases, 184 (12):1508–1517, 2001.

Robert J. Sampson, Jeffrey D. Monrenoff, and Thomas Gannon-Rowley. Assessing “neighborhood effects”: Social processes and new directions in research. Annual Review of Sociology, 28(1):443–478, 2002.

Thomas C. Schelling. Micromotives and Macrobehavior. Norton, 1978.

Avi Shmida and Stephan Ellner. Coexistence of plant species with similar niches. Vegetation, 58(1):29–55, 1984.

Rajesh Singh and Ronojoy Adhikari. Age-structured impact of social distancing on the COVID-19 epidemic in India. 2003.12005, 2020.

Se Jin Song, Christian Lauber, Elizabeth K Costello, Catherine A Lozupone, Gregory Humphrey, Donna Berg-Lyons, J Gregory Caporaso, Dan Knights, Jose C Clemente, Sara Nakielny, et al. Cohabiting family members share microbiota with one another and with their dogs. Elife, 2:e00458, 2013.

Hannes Svardal and Clau Rueffler. A general condition for adaptive genetic polymorphism in temporally and spatially heterogeneous environments. Theoretical Population Biology, 99:7–94, 2015.

Hannes Svardal, Claus Rueffler, and Joachim Hermisson. Comparing environmental and genetic variance as adaptive response to fluctuating selection. Evolution, 65(9):2493–2513, 2011.

Michael Turelli, Douglas W. Schemske, and Paulette Bierzychudek. Stable two-allele polymorphisms maintained by fluctuating fitnesses and seed banks: protecting the blues in Linanthus parryae. Evolution, 55 (7):1283–1298, 2001.

Robert Verity, Lucy C. Okell, Ilaria Dorigatti, et al. Estimates of the severity of COVID-19 disease. medRxiv, 2020.

Zunyou Wu and Jennifer M. McGoogan. Characteristics of and important lessons from the coronavirus disease 2019 (COVID-19) outbreak in China: Summary of a report of 72314 cases from the Chinese Center for Disease Control and Prevention. JAMA, 323(12):1239, 2020.

Hui-Ling Yen and Robert G. Webster. Pandemic Influenza as a Current Threat. Springer, 2009.

Juanjuan Zhang, Maria Litvinova, Yuxia Liang, Yan Wang, Wei Wang, Shanlu Zhao, Qianhui Wu, Stefano Merler, Cecile Viboud, Alessandro Vespignani, Marco Ajelli, and Hongjie Yu. Changes in contact patterns shape the dynamics of the COVID-19 outbreak in China. Science, 368(6498):1481–1486, 2020a.

Qian Zhang, Paul Bastard, Zhiyong Liu, et al. Inborn errors of type I IFN immunity in patients with life-threatening COVID-19. Science, 370(6515), 2020b.

Danping Zheng, Timur Liwinski, and Eran Elinav. Interaction between microbiota and immunity in health and disease. Cell Research, 30(6):492–506, 2020.

Xujuan Zhou, Enrico Coiera, Guy Tsafnat, Diana Arachi, Mei-Sing Ong, and Adam G. Dunn. Using social connection information to improve opinion mining: Identifying negative sentiment about HPV vaccines on Twitter. Medinfo, pages 761–765, 2015.

Daniel Zinder, Trevor Bedford, Sunetra Gupta, and Mercedes Pascual. The roles of competition and mutation in shaping antigenic and genetic diversity in influenza. PLOS Pathogens, 9(1):31003104, 2017a.

Daniel Zinder, Maria A. Riolo, Wood Robert J., and Mercedes Pascual. Role of competition in the strain structure of rotavirus under invasion and reassortment. bioRxiv, 2017b.

